# Humoral immune responses to the monovalent XBB.1.5-adapted BNT162b2 mRNA booster

**DOI:** 10.1101/2023.12.21.572575

**Authors:** Ulrika Marking, Oscar Bladh, Katherina Aguilera, Yiqiu Yang, Nina Greilert Norin, Kim Blom, Sophia Hober, Jonas Klingström, Sebastian Havervall, Mikael Åberg, Daniel J. Sheward, Charlotte Thålin

## Abstract

Continued SARS-CoV-2 evolution and immune escape necessitated the development of updated vaccines, and a monovalent vaccine incorporating the XBB.1.5 variant spike protein is currently being rolled out. Amidst the emergence of the highly mutated BA.2.86 lineage and against the backdrop of pronounced immune imprinting, it is important to characterize the antibody responses following vaccination, particularly in the elderly.

Here, we show that the monovalent XBB.1.5-adapted booster vaccination substantially enhanced both binding and neutralising antibody responses against a panel of variants, including BA.2.86, in an older population with four or more previous vaccine doses. Furthermore, neutralizing antibody titers to XBB.1.5 and BA.2.86 were boosted more strongly than titers to historical variants were.

Our findings thereby suggest increased vaccine induced protection against both antigenically matched variants, as well as the more distant BA.2.86 variant, and support current vaccine policies recommending a monovalent XBB.1.5 booster dose to older individuals.

## To the editor

SARS-CoV-2 viral evolution and immune evasion has reduced vaccine effectiveness. Bivalent booster doses delivering both the ancestral and BA.1/BA.5 spike fail to elicit strong omicron-specific humoral immune responses [1 2].

The global dominance of XBB sublineages prompted the development of further updated monovalent vaccines delivering the XBB.1.5 variant spike. Recently, however, a substantially mutated sublineage of BA.2, denoted BA.2.86, has emerged and several of its descendants are increasing in frequency. This raises questions whether the XBB.1.5-encoding vaccines will provide adequate protection against this emerging lineage. While preliminary data from phase 2/3 trials recently demonstrated a boost in neutralising antibody titres against both XBB lineages and BA.2.86 [3], ‘real-world’ data remains scarce [4].

Here, we characterised humoral responses in 24 individuals who were vaccinated with 30 µg of the monovalent XBB.1.5-adapted BNT162b2 mRNA booster vaccine in November 2023. Participants were enrolled through our ongoing observational Covid-19 Immunity (COMMUNITY) cohort study [5] with serological assessments at four-month intervals since April, 2020 (Appendix p2). Four participants contracted SARS-CoV-2 infections between XBB.1.5 vaccination and day 14 and thus were excluded. The majority of included participants (18/20) had received four or more vaccine doses prior to the monovalent XBB.1.5 booster, 18/20 had at least one confirmed prior SARS-CoV-2 infection, and the median age was 64 (IQR 59-67) (Supplementary table 1).

Using a pseudotyped virus-based neutralisation assay, we demonstrate a significant increase in neutralisation of all variants tested following the XBB.1.5 vaccine (Figure 1A). Geometric mean neutralising ID_50_ titres (GMT) to XBB.1.5 increased more than 10 times two weeks post vaccination (84 to 869). Importantly, neutralising titres to BA.2.86 were boosted with a similar magnitude (81 to 862) indicating that the XBB.1.5 vaccination elicits antibodies capable of neutralising BA.2.86. While wild type and BA.5 were still more potently neutralised than XBB.1.5 and BA.2.86, both before and after XBB.1.5 vaccination, the boost conferred by the vaccine to these historical variants was less pronounced (2.4- and 3.7-times increase in GMT at day 14) (Figure 1A). As a result, the difference between titres neutralizing wild type and omicron variants XBB.1.5 and BA.2.86, was also reduced by the XBB.1.5 vaccine (Supplementary Figure 2 A-B).

**Figure 1.**
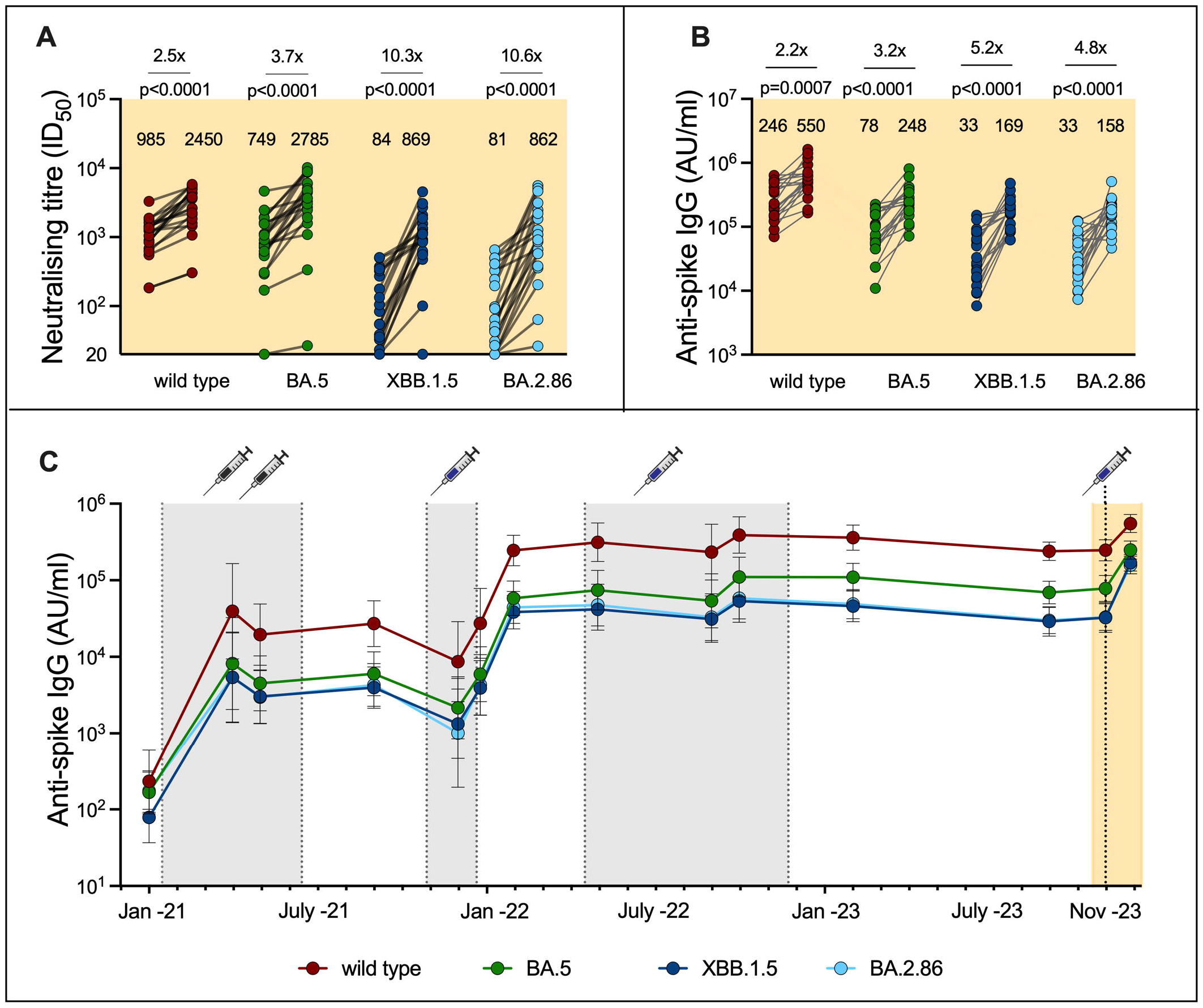
Monovalent XBB.1.5-adapted BNT162b2 booster vaccination elicits robust cross-neutralizing antibody responses. **(A)** Neutralization of wild type (B.1), BA.5, XBB.1.5 and BA.2.86 pseudoviruses by sera sampled 0-7 days prior to and 14 days after monovalent XBB.1.5-adapted BNT162b2 mRNA booster vaccination. The geometric mean ID_50_ titres are summarised above each group. ID_50_ titres below detection are plotted at the limit of detection (20). ID_50_ =half maximal inhibitory dilution. **(B)** Binding antibody titres (geometric mean titer) 0-7 days prior to and 14 days post XBB.1.5 vaccination. The geometric mean titres (in thousands) are summarized above each group. **(C)** Longitudinal binding antibody titres since primary vaccination in the cohort. The majority of participants had received four (12/20) or five (5/20) vaccine doses prior to the monovalent XBB.1.5-adapted booster vaccination. Gray shaded areas represent time intervals where primary and booster vaccinations were administered. The yellow shaded area represents the time interval between the sample taken prior to and 14 days post the monovalent XBB.1.5-adapted booster (i.e. the time period depicted in A and B).

Binding antibody titres to all variants tested also increased significantly following XBB.1.5 vaccination (Figure 1B and Supplementary Figure 1A), with relatively larger increases observed towards XBB.1.5 and BA.2.86 compared to wild type and BA.5 (Figure 1B). In line with neutralizing antibodies, there was a significant reduction in the ratio between wild type and XBB.1.5- and BA.2.86 binding titers. (Supplementary Figure 2 C-D). Longitudinal serological investigations conducted since April 2020, emphasize the noticeable reduction in the gap between antibody titres binding the wild type and titres binding omicron variants after the XBB.1.5 vaccination (Figure 1C and Supplementary Figure 1B).

In summary, monovalent XBB.1.5-adapted booster vaccination substantially enhanced both binding and neutralising antibody responses against a spectrum of variants in an older population with four or more previous vaccine doses. These data are largely consistent with a recent report from a younger population[4]. Against the backdrop of immunological imprinting and neutralizing antibody responses biased towards the ancestral variant, it is encouraging that the monovalent XBB.1.5-adapted booster elicits robust neutralizing antibody responses to XBB.1.5 and that recently circulating variants were boosted more strongly than responses to wild type and BA.5 variants. The significantly narrowed gap between titres to wild type and the XBB.1.5 and BA.2.86 variants post vaccination indicates that the monovalent XBB.1.5-adapted booster induces a shift in humoral responses toward the included variant, a feature the bivalent boosters appear to lack. Our findings thereby suggest increased vaccine induced protection against both antigenically matched variants, as well as the more distant BA.2.86 variant, and support current vaccine policies recommending a monovalent XBB.1.5 booster dose to older individuals.

## Acknowledgements

pCMV-dR8.2 dvpr was a gift from Bob Weinberg (Addgene plasmid # 8455; http://n2t.net/addgene:8455; RRID:Addgene_8455). pBOBI-FLuc was a gift from David Nemazee (Addgene plasmid # 170674; http://n2t.net/addgene:170674; RRID:Addgene_170674). We acknowledge the G2P-UK National Virology consortium funded by MRC/UKRI (grant ref: MR/W005611/1) and the Barclay Lab at Imperial College for providing the XBB.1.5 spike plasmid. We gratefully acknowledge all data contributors, i.e. the Authors and their Originating Laboratories responsible for obtaining the specimens, and their Submitting Laboratories that generated the genetic sequence and metadata and shared via the GISAID Initiative the data on which part of this research is based.

