## Supplementary Appendix for "Humoral immune responses to the monovalent XBB.1.5-adapted BNT162b2 mRNA booster"

^#^shared first authorship

*shared senior authorship

#### Table of contents

Methods	2

Study population	2

Antibody assays	2

Pseudovirus neutralization assay	2

Cell culture	2

Statistical analyses	3

Supplementary table and figures	4

Supplementary table 1	4

Supplementary figure 2	5

Supplementary figure 3-4	6

Supplementary references	7

### Methods

#### Study population

The COMMUNITY study investigates immune responses to SARS-CoV-2 infection and vaccination among 2,149 healthcare workers enrolled in April 2020 at Danderyd Hospital, Stockholm, Sweden [1-4]. Ongoing follow-ups every four months since enrolment include blood, saliva and nasal secretion sampling and collection of relevant clinical data. PCR screening sub studies are conducted during periods of high viral transmission. Vaccination details, including date and type of vaccine, are obtained from the Swedish vaccination registry (VAL Vaccinera), and data on positive SARS-CoV-2 PCR tests are sourced from the Swedish registry for communicable diseases (SmiNet). Clinical and demographic information, including immunocompromising conditions or treatments and information about positive rapid diagnostic tests (RDT), are collected via a smartphone application-based questionnaire during each follow-up visit.

Previous infection with SARS-CoV-2 is defined by one or more of the following criteria: seroconversion to the SARS-CoV-2 spike antigen before receiving primary vaccination, seroconversion to the SARS-CoV-2 nucleocapsid antigen, a positive SARS-CoV-2 PCR test registered in the national database of communicable diseases or in the study PCR screening programs or reporting of a positive result from an RDT at study follow-ups.

In this sub-study, we investigated serological responses to the monovalent XBB.1.5-adapted BNT162b2 mRNA booster dose (30 µg) in 24 participants. Blood samples were obtained up to 7 days prior to and 14 days post monovalent XBB.1.5-adapted BNT162b2 mRNA booster vaccination. Participants were tested with SARS-CoV-2 PCR at each sampling and encouraged to self-test with provided PCR kits if symptomatic during the study period. One participant tested positive with PCR on the day of XBB.1.5 vaccination and three participants tested positive 14 days post XBB.1.5 vaccination. These four participants were excluded from analysis of response to vaccination. The cohort is presented in table S1.

The study was approved by the Swedish Ethical Review Authority (dnr 2020-01653) and conducted in accordance with the declaration of Helsinki. Written informed consent was obtained from all study participants.

#### SARS-CoV-2 binding IgG

Serum IgG binding the SARS-CoV-2 wild type and variant (BA.5, XBB.1.5, BA.2.86, EG.5.1, FL.1.5.1, XBB.1.16, XBB.1.16.6, XBB.2.3) spike protein were analyzed with the V-PLEX SARS-CoV-2 panel 37 (Meso Scale Diagnostics, Maryland, USA) at dilutions 1:50000. Results are reported as arbitrary units (AU)/ml.

#### Cell culture

HEK293T cells (ATCC CRL-3216) and HEK293T cells stably expressing human ACE2 (HEK293T-ACE2) were cultured in Dulbecco’s Modified Eagle Medium (high glucose, with sodium pyruvate) that was supplemented with 10% fetal bovine serum, 100 units/ml Penicillin, and 100 μg/ml Streptomycin. Cultures were maintained in a humidified 37^o^C incubator (5% CO2).

#### Pseudovirus Neutralisation Assay

Pseudovirus neutralisation assays were performed as previously described [5] and production of the spike variants used here are described by Sheward et al [6]. Briefly, spike-pseudotyped lentivirus particles were produced from HEK293T cells by co-transfection of a spike-encoding plasmid, with a lentiviral gag-pol packaging plasmid (Addgene #8455), and a firefly luciferase transfer plasmid (Addgene #170674) using polyethylenimine. Pseudoviruses titrated to produce approximately 100,000 RLU were incubated with 8 serial 3-fold dilutions of serum (from 1:20) for 60 minutes at 37°C. Approximately 10,000 HEK293T-ACE2 cells were then added to each well, and plates were incubated at 37°C for 44-48 hours. Samples were tested against multiple variants ‘head-to-head’ using the same dilutions. Luminescence was measured on a GloMax Navigator Luminometer (Promega) using Bright-Glo luciferase substrate (Promega). Neutralisation was calculated relative to the mean of 8 control wells infected in the absence of antibody. Sera were heat inactivated at 56°C for 45 min prior to use in neutralisation assays. Reproducibility of measurements across two independent repeats are shown in Supplementary Figure 3.

#### Statistical analyses

ID_50_ values were calculated by fitting a four-parameter logistic curve and interpolating the dilution at which there is 50% neutralization. Continuous variables are presented as geometric mean titers (GMT) with corresponding 95% CI. Antibody titres before and after vaccination were compared using a Wilcoxon matched-pairs signed rank test. All fitting and statistical tests, including correlation analyses, were performed using Prism v10 (GraphPad Software, San Diego, California, USA).

### Supplementary Tables and Figures

|  | **Study participants (N=20)** |
| --- | --- |
| **Age** |  |
| Median [IQR] | 64.0 [59, 67] |
| **Sex** |  |
| Female | 17 (85%) |
| Male | 3 (15%) |
| **Time since last vaccine dose, days** |  |
| Median [IQR] | 381 [367, 401] |
| **Any prior SARS-CoV-2 infection** |  |
| No | 2 (10%) |
| Yes | 18 (90%) |
| **Infected prior to primary vaccine** |  |
| No | 13 (65%) |
| Yes | 7 (35%) |
| **Primary Vaccination Regimen** |  |
| ChAdOx1 x2 | 1 (5%) |
| ChAdOx1/BNT162b2 | 1 (5%) |
| BNT162b2 x2 | 18 (90%) |
| **Number of booster doses** |  |
| Median [IQR] | 2 [2, 3] |
| **Booster regimen** |  |
| Bivalent | 13 (65%) |
| Monovalent | 7 (35%) |

**Supplementary table 1.** Demographic characteristics and vaccination/infection histories of study participants (n=20). ChAdOx1; Vaxzevira (Covid-19 vaccine, AstraZeneca), BNT162b2; Comirnaty (Covid-19 vaccine, Pfizer), IQR; interquartile range.

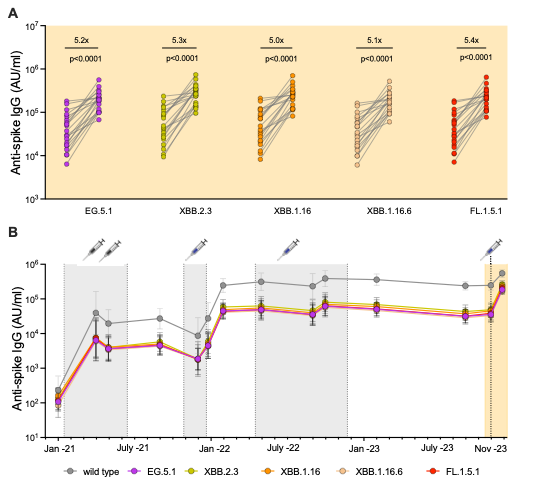

**Supplementary Figure 1.** Variant binding antibody titres **(A)** prior to and 14 days after monovalent XBB.1.5-adapted BNT162b2 mRNA booster vaccination and **(B)** longitudinally since primary vaccination in the cohort. The majority of participants had received four (12/20) or five (5/20) vaccine doses prior to the monovalent XBB.1.5-adapted booster vaccination. Gray shaded areas represent time intervals where primary and booster vaccination were administered. The yellow shaded area in B represents the time interval between the sample taken prior to and 14 days post the monovalent XBB.1.5-adapted booster (i.e. the time interval depicted in Figure A). Wild type binding titers are presented as a reference.

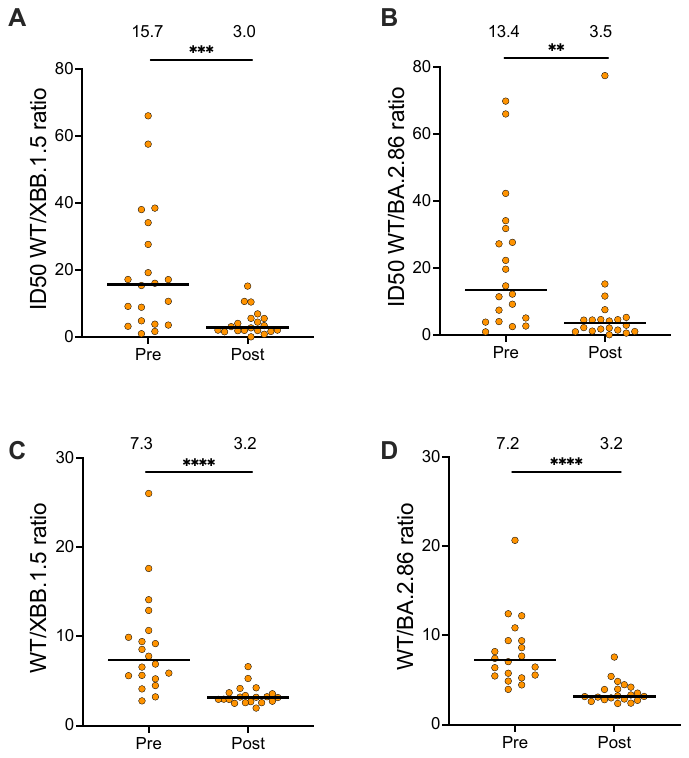

**Supplementary Figure 2.** Ratio of wild type/variant neutralizing (**A**-**B**) and binding (**C**-**D**) antibody titres before (pre) and after (post) the monovalent XBB.1.5-adapted BNT162b2 mRNA booster vaccination. Median ratios are summarised above each group. **** = p<0.0001, *** = p < 0.001, ** = p < 0.01. ID_50_ = half maximal inhibitory dilution, WT; wild type.

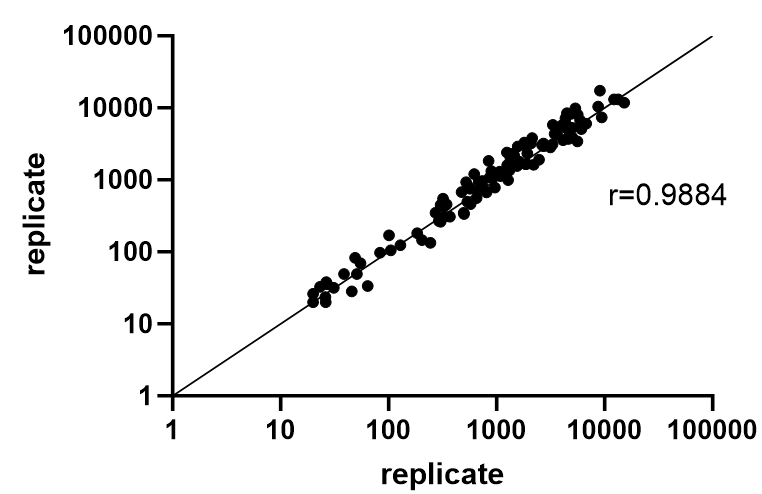

**Supplementary Figure 3. Reproducibility of neutralising titres measurements.** Shown are the titres for samples across two independent repeats overlaid onto the line of identity, and their log-domain Pearson correlation coefficient (r).
